# Death and rebirth of neural activity in sparse inhibitory networks

**DOI:** 10.1101/082974

**Authors:** David Angulo-Garcia, Stefano Luccioli, Simona Olmi, Alessandro Torcini

## Abstract

Inhibition is a key aspect of neural dynamics playing a fundamental role for the emergence of neural rhythms and the implementation of various information coding strategies. Inhibitory populations are present in several brain structures and the comprehension of their dynamics is strategical for the understanding of neural processing. In this paper, we clarify the mechanisms underlying a general phenomenon present in pulse-coupled heterogeneous inhibitory networks: inhibition can induce not only suppression of the neural activity, as expected, but it can also promote neural reactivation. In particular, for globally coupled systems, the number of firing neurons monotonically reduces upon increasing the strength of inhibition (neurons’ death). However, the random pruning of the connections is able to reverse the action of inhibition, i.e. in a sparse network a sufficiently strong synaptic strength can surprisingly promote, rather than depress, the activity of the neurons (neurons’ rebirth). Thus the number of firing neurons reveals a minimum at some intermediate synaptic strength. We show that this minimum signals a transition from a regime dominated by the neurons with higher firing activity to a phase where all neurons are effectively sub-threshold and their irregular firing is driven by current fluctuations. We explain the origin of the transition by deriving an analytic mean field formulation of the problem able to provide the fraction of active neurons as well as the first two moments of their firing statistics. The introduction of a synaptic time scale does not modify the main aspects of the reported phenomenon. However, for sufficiently slow synapses the transition becomes dramatic, the system passes from a perfectly regular evolution to an irregular bursting dynamics. In this latter regime the model provides predictions consistent with experimental findings for a specific class of neurons, namely the medium spiny neurons in the striatum.

## I. INTRODUCTION

The presence of inhibition in excitable systems induces a rich dynamical repertoire, which is extremely relevant for biological [13], physical [33] and chemical systems [82]. In particular, inhibitory coupling has been invoked to explain cell navigation [85], morphogenesis in animal coat pattern formation [46], and the rhythmic activity of central pattern generators in many biological systems [28, 45]. In brain circuits the role of inhibition is fundamental to balance massive recurrent excitation [72] in order to generate physiologically relevant cortical rhythms [12, 71].

Inhibitory networks are important not only for the emergence of rhythms in the brain, but also for the fundamental role they play in information encoding in the olfactory system [40] as well as in controlling and regulating motor and learning activity in the basal ganglia [5, 15, 47]. Furthermore, stimulus dependent sequential activation of neurons or group of neurons, reported for asymmetrically connected inhibitory cells [34, 52], has been suggested as a possible mechanism to explain sequential memory storage and feature binding [66].

These explain the long term interest for numerical and theoretical investigations of the dynamics of inhibitory networks. Already the study of globally coupled homogeneous systems revealed interesting dynamical features, ranging from full synchronization to clustering appearance [24, 81, 83], from the emergence of splay states [88] to oscillator death [6]. The introduction of disorder, e.g. random dilution, noise or other form of heterogeneity, in these systems leads to more complex dynamics, ranging from fast global oscillations [9] in neural networks and self-sustained activity in excitable systems [38], to irregular dynamics [3, 27, 30–32, 42, 49, 80, 88]. In particular, inhibitory spiking networks, due to *stable chaos* [62], can display extremely long erratic transients even in linearly stable regimes [3, 30, 31, 42, 49, 80, 87, 88].

One of the most studied inhibitory neural population is represented by medium spiny neurons (MSNs) in the striatum (which is the main input structure of the basal ganglia) [37, 56]. In a series of papers Ponzi and Wickens have shown that the main features of the MSN dynamics can be reproduced by considering a sparsely connected inhibitory network of conductance based neurons subject to external stochastic excitatory inputs [63–65]. Our study has been motivated by an interesting phenomenon reported for this model in [65]: namely, upon increasing the synaptic strength the system passes from a regularly firing regime, characterized by a large part of quiescent neurons, to a biologically relevant regime where almost all cells exhibit a bursting activity, characterized by an alternation of periods of silence and of high firing. The same phenomenology has been recently reproduced by employing a much simpler neural model [1]. Thus suggesting that this behaviour is not related to the specific model employed, but it is indeed a quite general property of inhibitory networks. However, it is still unclear the origin of the phenomenon and the minimal ingredients required to observe the emergence of this effect.

In order to exemplify the problem addressed in this paper we report in Fig. 1 the fraction of active neurons *n*_*A*_(i.e. the ones emitting at least one spike during the simulation time) as a function of the strength of the synaptic inhibition *g* in an heterogenous network. For a fully coupled network, *n*_*A*_ has a monotonic decrease with *g* (Fig. 1 (a)), while for a sparse network *n*_*A*_ has a non monotonic behaviour, displaying a minimum at an intermediate strength *g*_*m*_ (Fig. 1 (b)). In fully coupled networks the effect of inhibition is simply to reduce the number of active neurons (*neurons’ death*). However, quite counter-intuitively, in presence of dilution by increasing the synaptic strength the previously silenced neurons can return to fire (*neurons’ rebirth*). Our aim is to clarify the physical mechanisms underlying neuron’s death and rebirth, which are at the origin of the behaviour reported in [1, 65].

**FIG. 1.**
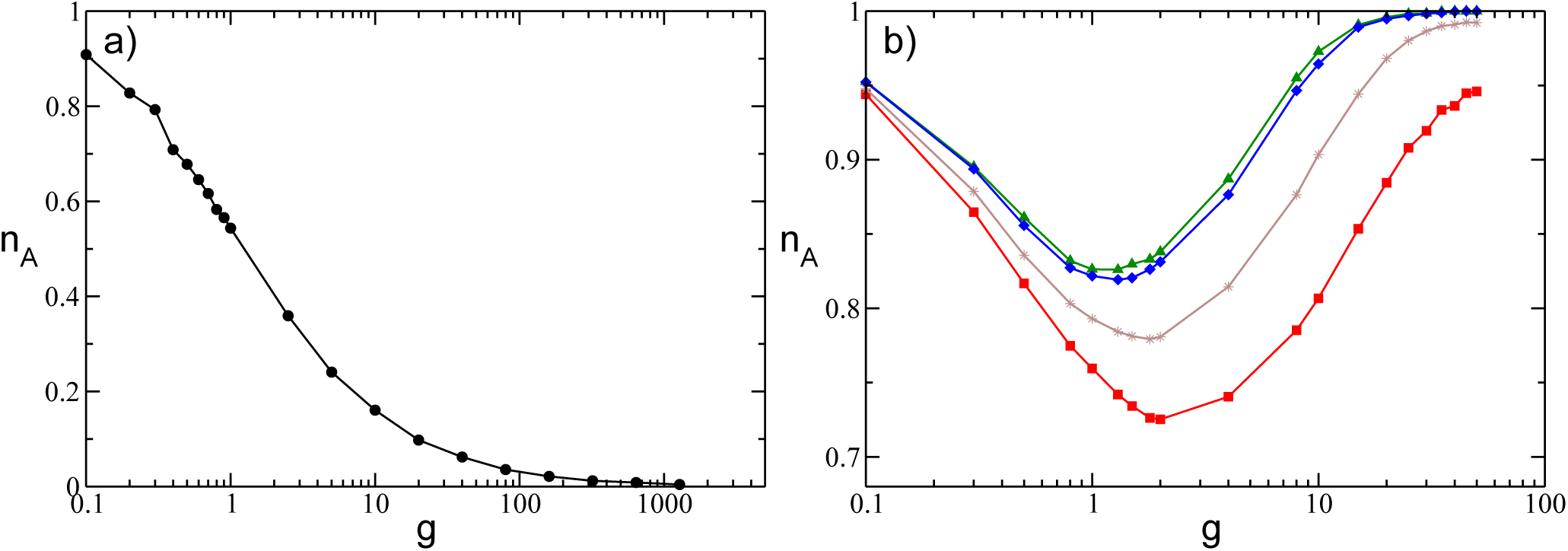
Fraction of active neurons *n*_*A*_ as a function of the inhibitory synaptic strength *g* for a globally coupled system (a), where *K = N* − 1, and a sparsely connected network with *K* = 20 (b). In a) is reported the asymptotic value *n*_*A*_ calculated after a time *t*_*s*_ = 1 × 10^6^. Conversely in b), *n*_*A*_ is reported at successive times: namely, *t*_*s*_ = 985 (red squares), *t*_*s*_ = 1.1 × 10^4^ (brown stars), *t*_*s*_ = 5 × 10^5^ (blue diamonds) and *t*_*s*_ = 1 × 10^6^ (green triangles). These values have been also averaged over 10 random realizations of the network. The reported data refer to instantaneous synapses, to a system size *N* = 400 and an uniform distribution *P(I)* with [*l*_1_,: *l*_2_] = [1.0 : 1.5] and *θ* = 1.

In particular, we consider a deterministic network of purely inhibitory pulse-coupled Leaky Integrate-and-Fire (LIF) neurons with an heterogeneous distribution of excitatory DC currents, accounting for the different level of excitability of the neurons. The evolution of this model is studied for fully coupled and sparse topology as well as for synapses with different time courses. For the fully coupled case, it is possible to derive, within a self-consistent mean field approach, the analytic expressions for the fraction of active neurons and of the average firing frequency 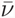 as a function of the coupling strength *g.* In this case the monotonic decrease of *n*_*A*_ with *g* can be interpreted as a *Winner Takes All* (WTA) mechanism [16, 21, 86], where only the the most excitable neurons survive to the inhibition increase. For sparse networks, the neurons’ rebirth can be interpreted as a reactivation process induced by erratic fluctuations in the synaptic currents. Within this framework it is possible to obtain analytically for instantaneous synapses, a closed set of equations for *n*_*A*_ as well as for the average firing rate and coefficient of variation as a function of the coupling strength. In particular, the firing statistics of the network can be obtained via a mean-field approach by extending the formulation derived in [69] to account for synaptic shot noise with constant amplitude. The introduction of a finite synaptic time scale do not modify the overall scenario as far as this is shorter than the membrane time constant. As soon as the synaptic dynamics becomes slower, the phenomenology of the transition is modified. At *g* < *g*_*m*_ we have a *frozen phase* where *n*_*A*_ does not evolve in time on the explored time scales, since the current fluctuations are negligible. Above *g*_*m*_ we have a bursting regime, which can be related to the emergence of correlated fluctuations as discussed within the adiabatic approach in [50, 51].

The paper is organized as follows: In Sect. II we present the models that will be considered in the paper as well as the methods adopted to characterize its dynamics. In Sect. III we consider the globally coupled network where we provide self-consistent expressions accounting for the fraction of active neurons and the average firing rate. Section IV is devoted to the study of sparsely connected networks with instantaneous synapses and to the analytic derivation of the set of self-consistent equations providing *n*_*A*_, the average firing rate and the coefficient of variation. In section V we discuss the effect of synaptic filtering with a particular attention on slow synapses. Finally in Sect. VI we briefly discuss the obtained results with a focus on the biological relevance of our model.

## II. MODEL AND METHODS

We examine the dynamical properties of an heterogeneous inhibitory sparse network made of *N* LIF neurons. The time evolution of the membrane potential *v*_*i*_ of the *i*-th neuron is ruled by the following first order ordinary differential equation:

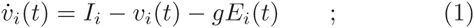

where *g* > 0 is the inhibitory synaptic strength, *I*_*i*_ is the neuronal excitability of the *i*-th neuron encompassing both the intrinsic neuronal properties and the excitatory stimuli originating from areas outside the considered neural circuit and *E*_*i*_(*t*) represents the synaptic current due to the recurrent interactions within the considered network. The membrane potential *v*_*i*_ of neuron *i* evolves accordingly to Eq. (1) until it overcomes a constant threshold *θ* = 1, this leads to the emission of a spike (action potential) transmitted to all the connected post-synaptic neurons, while *v*_*i*_ is reset to its rest value *v*_*r*_ = 0. The model in (1) is expressed in adimensional units, this amounts to assume a membrane time constant *τ*_*m*_ = 1, for the conversion to dimensional variables see Appendix A. The heterogeneity is introduced in the model by assigning to each neuron a different value of input excitability *I*_*i*_ drawn from a flat distribution *P*(*I*), whose support is *I* ∈ [*l*_1_ : *l*_2_] with *l*_1_ ≥ *θ*, therefore all the neurons are supra-threshold.

The synaptic current *E*_*i*_(*t*) is given by the linear superposition of all the inhibitory post-synaptic potentials (IPSPs) *η*(*t*) emitted at previous times 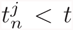 by the presynaptic neurons connected to neuron *i,* namely

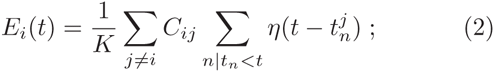

where *K* is the number of pre-synaptic neurons. *C*_*ij*_ represent the elements of the *N* × *N* connectivity matrix associated to an undirected random network, whose entries are 1 if there is a synaptic connection from neuron *j* to neuron *i,* and 0 otherwise. For the sparse network, we select randomly the matrix entries, however to reduce the sources of variability in the network, we assume that the number of pre-synaptic neurons is fixed, namely ∑_*j*≠*i*_*C*_*ij*_ = *K* << *N* for each neuron *i,* where autaptic connections are not allowed. We have verified that the results do not change if we choose randomly the links accordingly to an Erdös-Renyi distribution with a probability *K/N.* For a fully coupled network we have *K = N* − 1.

The shape of the IPSP characterize the type of filtering performed by the synapses on the received action potentials. We have considered two kind of synapses, instantaneous ones, where *η*(*t*) = *δ*(*t*), and synapses where the PSP is an *α*-pulse, namely

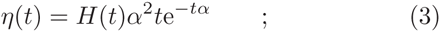

with *H* denoting the Heaviside step function. In this latter case the rise and decay time of the pulse are the same, namely *τ*_*α*_ = 1/*α,* and therefore the pulse duration *τ*_*P*_ can be assumed to be twice the characteristic time *τ*_*α*_. The equations of the model Eqs. (1) and (2) are integrated exactly in terms of the associated *event driven maps* for different synaptic filtering, these correspond to Poincaré maps performed at the firing times (for details see Appendix A) [53, 89].

For instantaneous synapses, we have usually considered system sizes *N* = 400 and *N* = 1,400 and for the sparse case in-degrees 20 ≤ *K* ≤ 80 for *N* = 400 and 20 ≤ *K* ≤ 600 for *N* = 1400 with integration times up to *t*_*s*_ = 1 × 10^6^. For synapses with a finite decay time we limit the analysis to *N* = 400 and *K* = 20 and to maximal integration times *t*_*s*_ = 1 × 10^5^. Finite size dependences on *N* are negligible with these parameter choices, as we have verified.

In order to characterize the network dynamics we measure the fraction of active neurons *n*_*A*_(*t*_*S*_) at time *t*_*S*_, i.e. the fraction of neurons emitting at least one spike in the time interval [0 : *t*_*S*_]. Therefore a neuron will be considered silent if it has a frequency smaller than 1/*t*_*S*_, with our choices of *t*_*S*_ = 10^5^ – 10^6^, this corresponds to neurons with frequencies smaller than 10^−3^ – 10^−4^ Hz, by assuming as timescale a membrane time constant *τ*_*m*_ = 10 ms. The estimation of the number of active neurons is always started after a sufficiently long transient time has been discarded, usually corresponding to the time needed to deliver 10^6^ spikes in the network.

Furthermore, for each neuron we estimate the time averaged inter-spike interval (ISI) *T*_*ISI*_, the associated firing frequency *ν* = 1/*T*_*ISI*_, as well as the coefficient of variation *CV,* which is the ratio of the standard deviation of the ISI distribution divided by *T*_*ISI*_. For a regular spike train *CV* = 0, for a Poissonian distributed one *CV* = 1, while *CV* > 1 is usually an indication of bursting activity. The indicators usually reported in the following to characterize the network activity are ensemble averages over all the active neurons, which we denote as 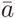 for a generic observable *a*.

To analyze the linear stability of the dynamical evolution we measure the maximal Lyapunov exponent *λ*, which is positive for chaotic evolution, and negative (zero) for stable (marginally stable) dynamics [4]. In particular, by following [2, 55] *λ* is estimated by linearizing the corresponding event driven map.

## III. FULLY COUPLED NETWORKS: WINNER TAKES ALL

In the fully coupled case we observe that the number of active neurons *n*_*A*_ saturates, after a short transient, to a value which remains constant in time. In this case, it is possible to derive a self-consistent mean field approach to obtain an analytic expression for the fraction of active neurons *n*_*A*_ and for the average firing frequency 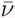 of the neurons in the network. In a fully coupled network each neuron receives the spikes emitted by the other *K = N*−1 neurons, therefore each neuron is essentially subject to the same effective input *µ,* apart corrections *O*(1/*N*).

The effective input current, for a neuron with an excitability *I,* is given by

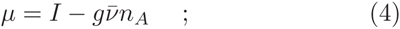

where *n*_*A*_(*N −* 1) is the number of active pre-synaptic neurons assumed to fire with the same average frequency 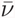.

In a mean field approach, each neuron can be seen as isolated from the network and driven by the effective input current *µ.* Taking into account the distribution of the excitabilities *P*(*I*) one obtains the following self-consistent expression for the average firing frequency

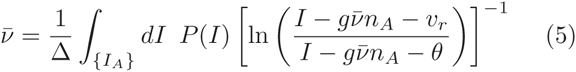

where the integral is restricted only to active neurons, i.e. to *I* ∈ {*I*_*A*_} values for which the logarithm is defined, while ∆ = ∫_{*I*_*A*_}_ *dI P*(*I*) is the measure of their support. In (5) we have used the fact that for an isolated LIF neuron with constant excitability *C,* the ISI is simply given by *T*_*ISI*_ = ln[(*C* − *v*_*r*_)/(*C − θ*)] [11].

An implicit expression for *n*_*A*_ can be obtained by estimating the neurons with effective input *µ* > *θ,* in particular the number of silent neurons is given by

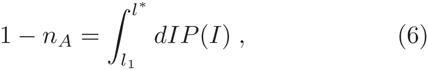

where *l*_1_ is the lower limit of the support of the distribution, while 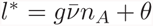. By solving self-consistently Eqs.(5) and (6) one can obtain the analytic expression for *n*_*A*_ and 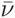 for any distribution *P*(*I*).

In particular, for excitabilities distributed uniformly in the interval [*l*_1_ : *l*_2_], the expression for the average frequency Eq. (5) becomes

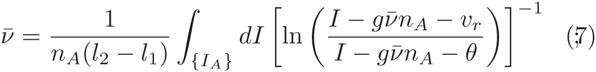

while the number of active neurons is given by the following expression

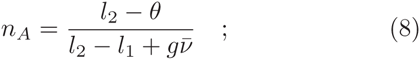

with the constraint that *n*_*A*_ cannot be larger than one.

The analytic results for these quantities compare quite well with the numerical findings estimated for different distribution intervals [*l*_1_ : *l*_2_], different coupling strengths and system sizes, as shown in Fig. 2. Apart for definitely large coupling *g* > 10 where some discrepancies among the mean field estimations and the simulation results for 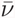 is observable (see Fig. 2 (b)). These differences are probably due to the discreteness of the pulses, which cannot be neglected for very large synaptic strengths.

**FIG. 2.**
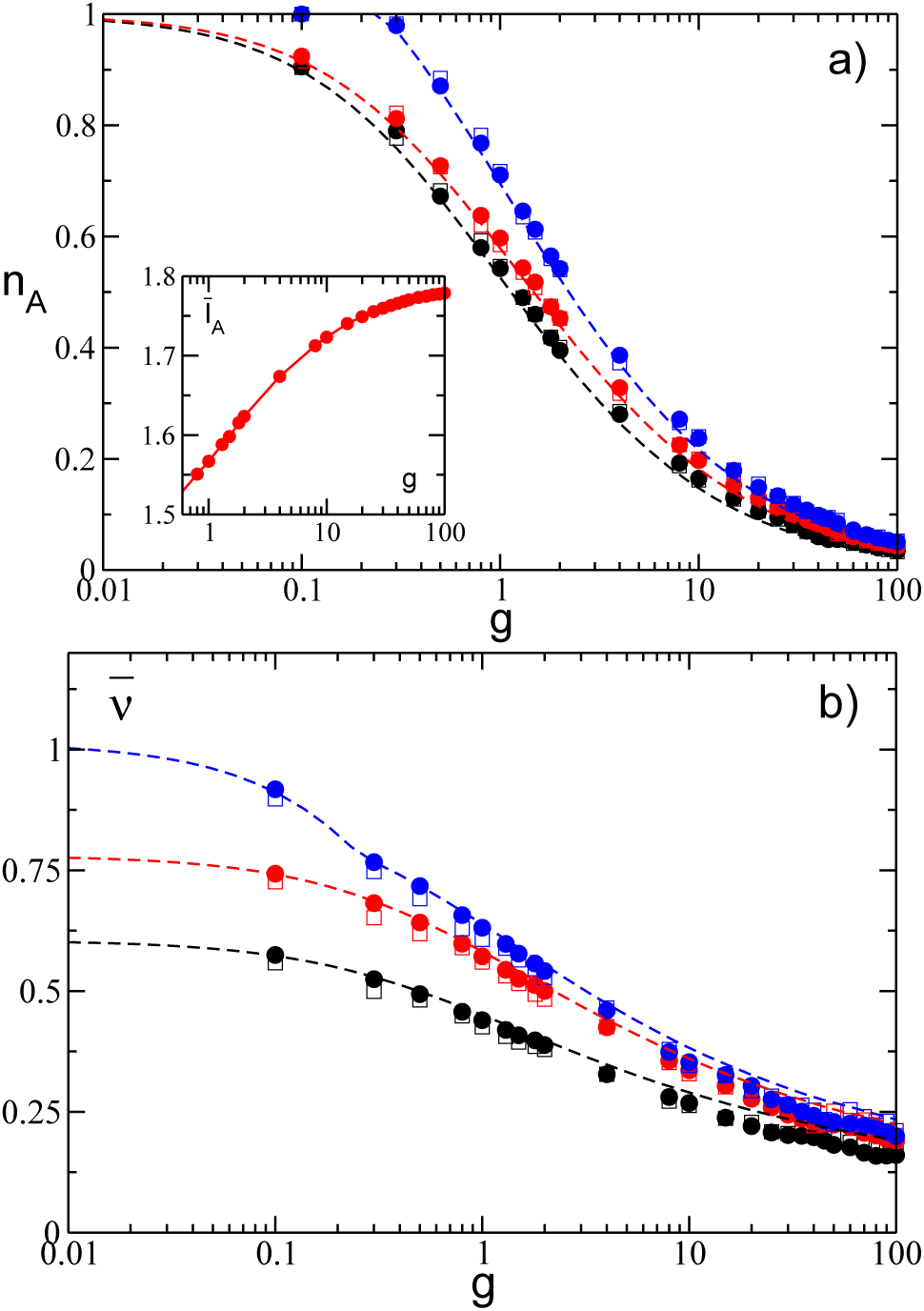
Fraction of active neurons *n*_*A*_ (a) and average network’s frequency 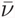 (b) as a function of the synaptic strength *g* for uniform distributions *P*(*I*) with different supports. Inset: average neuronal excitability of the active neurons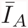 versus *g*. Empty (filled) symbols refer to numerical simulation with *N* = 400 (*N* = 1400) and dashed lines to the corresponding analytic solution. Symbols and lines correspond from bottom to top to [*l*_1_ : *l*_2_] = [1.0 : 1.5] (black); [*l*_1_ : *l*_2_] = [1.0 : 1.8] (red) and [*l*_1_ : *l*_2_] = [1.2 : 2.0] (blue). The data have been averaged over a time interval *t*_*S*_ = 1 × 10^6^ after discarding a transient of 10^6^ spikes.

As a general feature we observe that *n*_*A*_ is steadily decreasing with *g*, thus indicating that a group of neurons with higher effective inputs (*winners*) silence the other neurons (*losers*) and that the number of *winners* eventually vanishes for sufficiently large coupling in the limit of large system sizes. Furthermore, the average excitability of the active neurons (the *winners*) 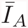 increases with *g*, as shown in the inset of Fig. 2 (a), thus revealing that only the neurons with higher excitabilities survive to the silencing action exerted by the other neurons. At the same time, as an effect of the growing inhibition the average firing rate of the *winners* dramatically slows down. Therefore despite the increase of 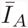 the average effective input 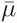 indeed decreases for increasing inhibition. This represents a clear example of the winner takes all (WTA) mechanism obtained via (lateral) inhibition, which has been shown to have biological relevance for neural systems [20, 22, 59, 86].

It is important to understand which is the minimal coupling value *g*_*c*_ for which the firing neurons start to die. In order to estimate *g*_*c*_ it is sufficient to set *n*_*A*_ = 1 in Eqs. (7) and (8). In particular, one gets

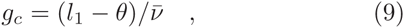

thus for *l*_1_ = *θ* even an infinitesimally small coupling is in principle sufficient to silence some neuron. Furthermore, from Fig. 3 (a) it is evident that whenever the excitabilities become homogeneous, i.e. for *l*_1_ → *l*_2_, the critical synaptic coupling *g*_*c*_ diverges towards infinite. Thus heterogeneity in the excitability distribution is a necessary condition in order to observe a (gradual) neurons’ death, as shown in Fig. 2 (a).

**FIG. 3.**
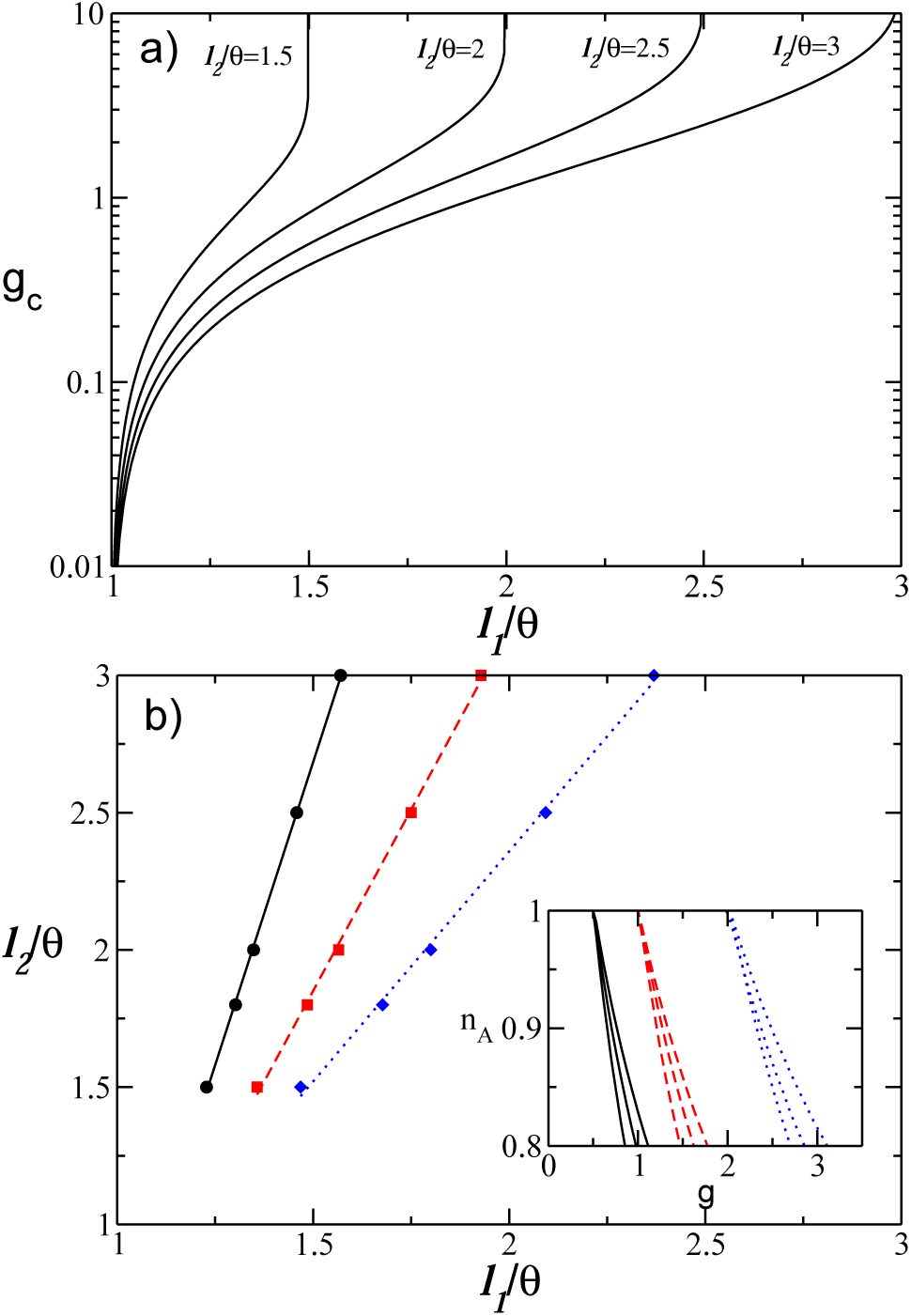
a) Critical value *g*_*c*_ as a function of the lower value of the excitability *l*_1_ for several choices of the upper limit *l*_2_. b) Isolines corresponding to constant values of *g*_*c*_ in the (*l*_1_, *l*_2_)-plane: namely, *g*_*c*_ = 0.5 (black solid line), *g*_*c*_ = 1.0 (red dashed line), *g*_*c*_ = 2.0 (blue dotted line). Inset: Dependence of *n*_*A*_ on *g* for three couples of values (*l*_1_, *l*_2_) chosen along each of the isolines reported in the main figure.

This is in agreement with the results reported in [7], where homogeneous fully coupled networks of inhibitory LIF neurons have been examined. In particular, for finite systems and slow synapses the authors in [7] reveal the existence of a sub-critical Hopf bifurcation from a fully synchronized state to a regime characterized by oscillator death occurring at some critical *g*_*c*_. However, in the thermodynamic limit *g*_*c*_ → ∞ for fast as well as slow synapses, in agreement with our mean field result for instantaneous synapses.

We also proceed to investigate the isolines corresponding to the same critical *g*_*c*_ in the (*l*_1_, *l*_2_)-plane, the results are reported in Fig. 3 (b) for three selected values of *g*_*c*_. It is evident that the *l*_1_ and *l*_2_-values associated to the isolines display a direct proportionality among them. However, despite lying on the same *g*_*c*_-isoline, different parameter values induce a completely different behaviour of *n*_*A*_ as a function of the synaptic strength, as shown in the inset of Fig. 3 (b).

Direct simulations of the network at finite sizes, namely for *N* = 400 and *N* = 1400, show that for sufficiently large coupling neurons with similar excitabilities tend to form clusters, similarly to what reported in [42], for the same model here studied, but with a delayed pulse transmission. However, at variance with [42], the overall macroscopic dynamics is asynchronous and no collective oscillations can be detected for the whole range of considered synaptic strengths.

## IV. SPARSE NETWORKS : NEURONS’ REBIRTH

In this Section we will consider a network with sparse connectivity, namely each neuron is supra-threshold and it receives instantaneous IPSPs from *K* << *N* randomly chosen neurons in the network. Due to the sparseness, the input spike trains can be considered as uncorrelated and at a first approximation it can be assumed that each spike train is Poissonian with a frequency 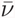 correspondent to the average firing rate of the neurons in the network [8, 9]. Usually, the mean activity of a LIF neural network has been estimated in the context of the diffusion approximation [68, 78]. This approximation is valid whenever the arrival frequency of the IPSPs is high with respect to the firing emission, while the amplitude of each IPSPs (namely, *G = g/K*) is small with respect to the firing threshold *θ.* This latter hypothesis in our case is not valid for sufficiently large (small) synaptic strength *g* (in-degree *K*), therefore the synaptic inputs should be treated as shot noise. In particular, here we apply an extended version of the approach derived by Richardson and Swabrick in [69] to estimate the average firing rate and the average coefficient of variation for LIF neurons subject to inhibitory shot noise with constant amplitude.

At variance with the fully coupled case, the fraction of active neurons *n*_*A*_ does not saturate to a constant value for sufficiently short times. Instead, *n*_*A*_ increases in time, due to the rebirth of losers which have been previously silenced by the firing activity of the winners. As shown in in Fig. 1(b), the variation of *n*_*A*_ slows down for increasing integration times *t*_*S*_ and *n*_*A*_ approaches some apparently asymptotic profile for sufficiently long integration times. Furthermore, *n*_*A*_ has a non monotonic behaviour with *g*, as opposite to the fully coupled case. In particular, *n*_*A*_ reveals a minimum *n*_*A*_*m*__ at some intermediate synaptic strength *g*_*m*_ followed by an increase towards *n*_*A*_ = 1 at large *g.* As we have verified, as far as 1 < *K* << *N* finite size effects are negligible and the actual value of *n*_*A*_ depends only on the in-degree *K* and the considered simulation time *t*_*S*_. In the following we will try to explain the origin of such a behaviour.

Despite the model is fully deterministic, due to the random connectivity the observed phenomena can be interpreted in the framework of activation processes. In particular, we can assume that each neuron in the network will receive *n*_*A*_*K* independent Poissonian trains of inhibitory kicks of constant amplitude *G* characterized by an average frequency 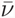, thus each synaptic input can be regarded as a single Poissonian train with total frequency 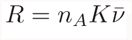, Therefore, each neuron, characterized by its own excitability *I*, will be subject to an average effective input *µ*(*I)* (as reported in Eq. (4)) plus fluctuations in the synaptic current of amplitude

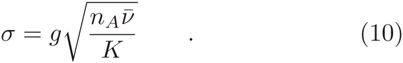

Indeed, we have verified that (10) gives a quantitatively correct estimation of the synaptic current fluctuations over the whole range of synaptic coupling here considered (as shown in Fig. 4). For instantaneous IPSP, these fluctuations are due to stable chaos [62], since the maximal Lyapunov exponent is negative for the whole range of coupling, as we have verified. Therefore, as reported by many authors, erratic fluctuations in inhibitory neural networks with instantaneous synapses are due to finite amplitude instabilities, while at the infinitesimal level the system is stable [3, 30, 31, 42, 49, 80, 88].

**FIG. 4.**
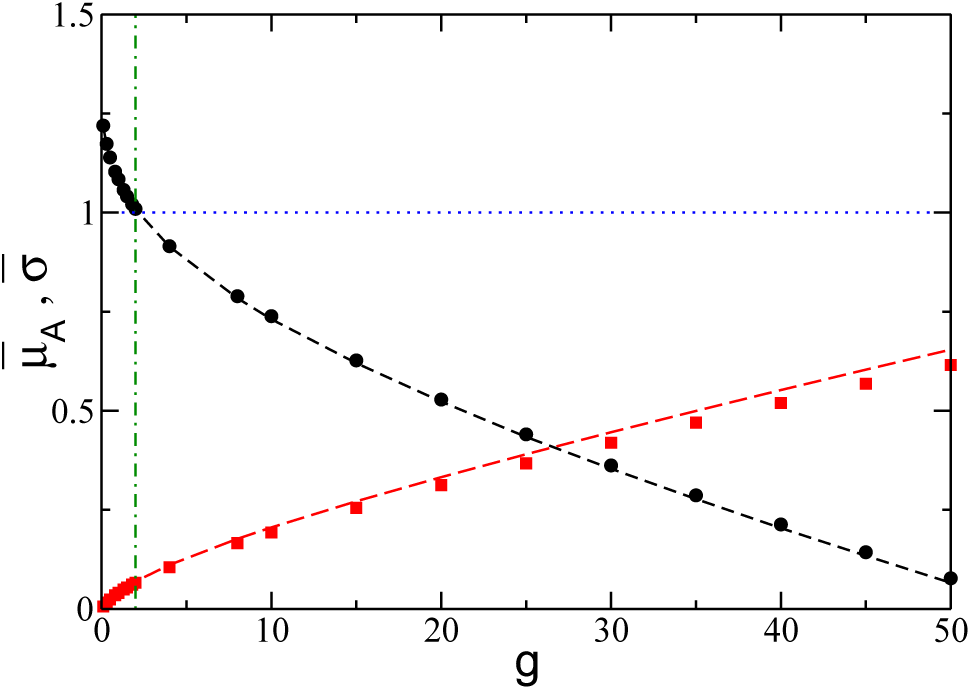
Effective average input of the active neurons 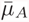 (black circles) and average fluctuations of the synaptic currents 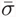 (red squares) as a function of the inhibitory coupling *g.* The threshold potential *θ* = 1 is marked by the (blue) horizontal dotted line and *g*_*m*_ by the (green) vertical dash-dotted line. The dashed black (red) line refer to the theoretical estimation for *µ*_*A*_ (σ) reported in Eq. (4) (Eq. (10)) and averaged only over the active neurons. The data refer to N = 1400, *K* = 140, [*l*_1_ : *l*_2_] = [1.0 : 1.5] and to a simulation time *t*_*s*_ = 1 × 10^6^.

In this picture, the silent neurons stay in a quiescent state corresponding to the minimum of the effective potential 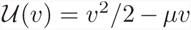 and in order to fire they should overcome a barrier 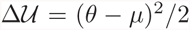. The average time *t*_*A*_ required to overcome such barrier can be estimated accordingly to the Kramers’ theory for activation processes [26, 78], namely

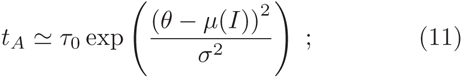

where *τ*_0_ is an opportune time scale.

It is reasonable to assume that at a given time *t*_*S*_ all neurons with *t*_*A*_ < *t*_*S*_ will have fired at least once, and the fraction of active neurons *n*_*A*_ at that time can be estimated by identifying the specific neuron for which *t*_*S*_ = *t*_*A*_. This neuron will be characterized by the following excitability

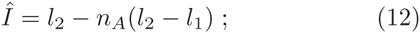

for excitabilities *I* uniformly distributed in the interval [*l*_1_ : *l*_2_]. In order to obtain the fraction of active neurons at time *t*_*S*_, one should solve the equation (11) for the neuron with excitability 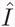 by setting *t*_*S*_ = *t*_*A*_, thus obtaining the following solution

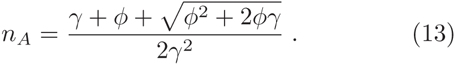

where

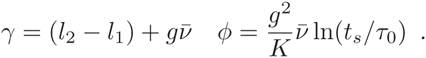

Equation (13) gives the dependence of *n*_*A*_ on the coupling strength *g* for a fixed integration time *t*_*S*_, whenever we can provide the value of the average frequency 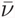. This in turn can be obtained analytically by following the approach introduced in [69] for LIF neurons subject to excitatory and inhibitory synaptic shot noise with exponentially distributed PSP amplitudes. In Appendix B we detail how to obtain with this method 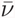 for inhibitory spike trains with constant PSP amplitudes. In particular, the average frequency can be found via the following self consistent equation

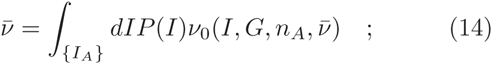

where the explicit expression of *ν*_0_ is given by Eq. (31) in Appendix B.

The simultaneous solution of Eqs. (13) and (14) provide a theoretical estimation of *n*_*A*_ and 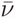 for the whole considered range of synaptic strength, once the unknown time scale *τ*_0_ is fixed. This time scale has been determined via an optimal fitting procedure for sparse networks with *N* = 400 and *K* = 20, 40 and 80 at a fixed integration time *t*_*S*_ = 1 × 10^6^. The results for *n*_*A*_ are reported in Fig. 5 (a), the estimated curves reproduce reasonably well the numerical data for *K* = 20 and 40, while for *K* = 80 the agreement worsens at large coupling strengths. This should be due to the fact that by increasing *g* and *K* the spike trains stimulating each neuron cannot be assumed to be completely independent, as done for the derivation of Eqs. (13) and (14). Nevertheless, the average frequency 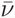 is quantitatively well reproduced for the considered *K* values over the entire range of the synaptic strengths, as it is evident from Figs. 5 (b-d).

**FIG. 5.**
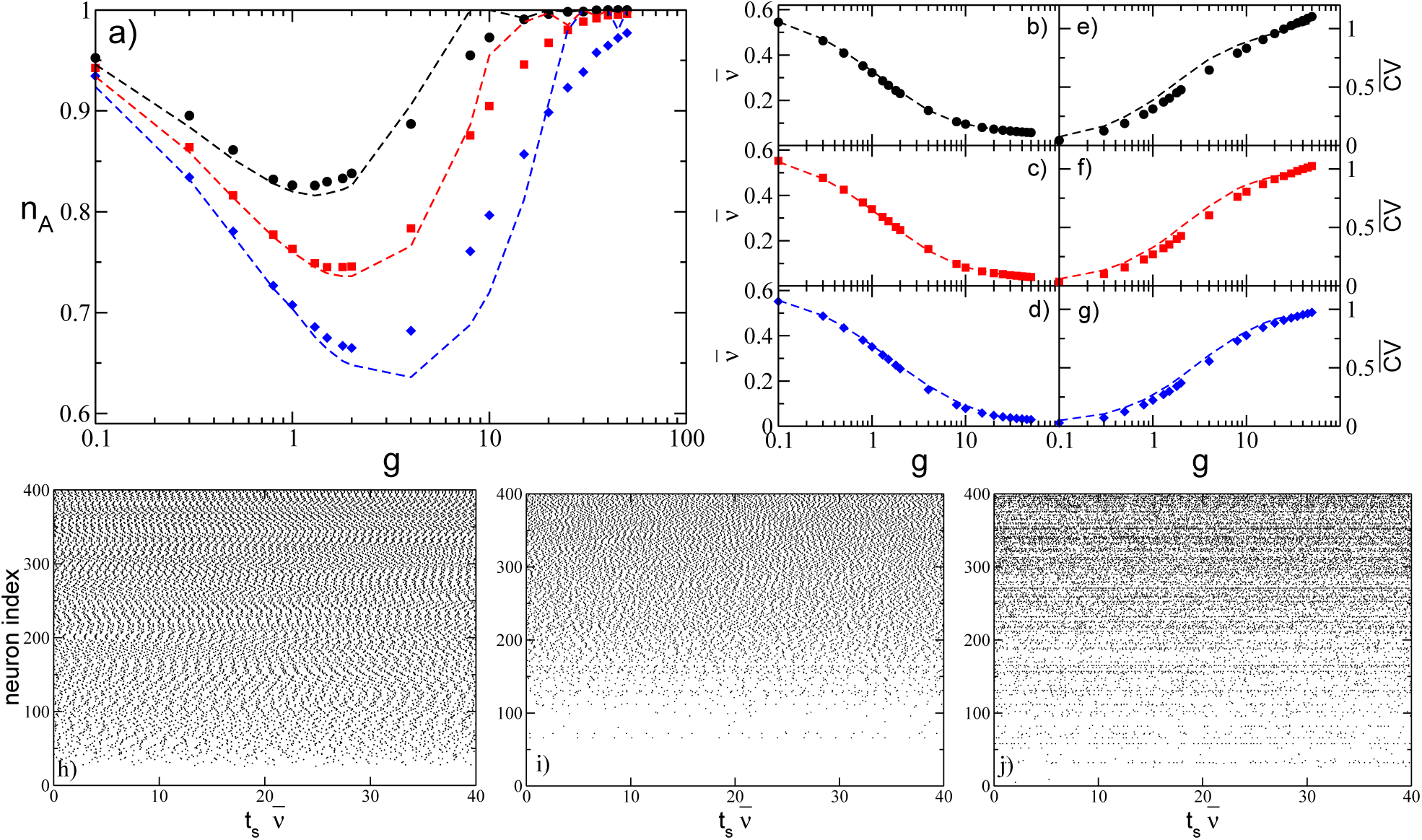
a) Fraction of active neurons *n*_*A*_ as a function of inhibition for several values of *K.* b-d) Average network’s firing rate 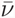 for the same cases depicted in a), and the corresponding 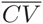 (e-f). In all panels, filled symbols correspond to numerical data and dashed lines to analytic values: black circles correspond to *K* = 20 (*τ*_0_ = 11), red squares to *K* = 40 (*τ*_0_ = 19) and blue diamonds to *K* = 80 (*τ*_0_ = 26.6). The data are averaged over a time interval *t*_*s*_ = 1 × 10^6^ and 10 different realizations of the random network. h-j) Raster plots for three different synaptic strenghts for *N* = 400 and *K* = 20: namely, h) *g* = 0.1; i) *g* = 1 and *j*) *g* = 10. The corresponding value for the fractionof active neurons, average frequency and average coefficient of variation are *n*_*A*_ = (0.95,0.83,0.97), 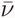= (0.55,0.32,0.10) and 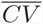= (0.04,0.31,0.83), respectively. The neurons are ordered in terms of their intrinsic excitability and the time is rescaled by the average frequency 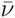. The remaining parameters as in Fig. 1 (b).

We have also estimated analytically the average coefficient of variation of the firing neurons 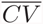 by extending the method derived in [69] to obtain the response of a neuron receiving synaptic shot noise inputs. The analytic expressions of the coefficient of variation for LIF neurons subject to inhibitory shot noise with fixed post-synaptic amplitude are obtained by estimating the second moment of the associated first-passage-time distribution, the details are reported in Appendix C. The coefficient of variation can be estimated, once the self consistent values for *n*_*A*_ and 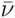 have been obtained via Eqs. (13) and (14). The comparison with the numerical data, reported in Figs 5 (e-g), reveals a good agreement over the whole range of synaptic strengths for all the considered in-degrees.

At sufficiently small synaptic coupling the neurons fire tonically and almost independently, as clear from the raster plot in Fig 5 (h) and by the fact that 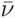 approaches the average value for the uncoupled system (namely 0.605) and 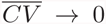. By increasing the coupling, *n*_*A*_ decreases, as an effect of the inhibition more and more neurons are silenced and the average firing rate decrease, at the same time the dynamics becomes slightly more irregular as shown in Fig 5 (i). At large coupling *g* > *g*_*m*_, a new regime appears, where almost all neurons become active but with an extremely slow dynamics which is essentially stochastic with 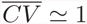, as testified also by the raster plot reported in Fig 5 (j).

Furthermore, from Fig. 5(a) it is clear that the value of the minimum of the fraction of active neurons *n*_*Am*_ decreases by increasing the network in-degree *K*, while *g*_*m*_ increases with *K.* This behaviour is further investigated in a larger network, namely *N* = 1400, and reported in the inset of Fig. 6. It is evident that *n*_*A*_ stays close to the globally coupled solutions over larger and larger intervals for increasing *K.* This can be qualitatively understood by the fact that the current fluctuations Eq. (10), responsible for the rebirth of silent neurons, are proportional to *g* and scales as 1/*√K*, therefore at larger in-degree the fluctuations have similar amplitudes only for larger synaptic coupling.

**FIG. 6.**
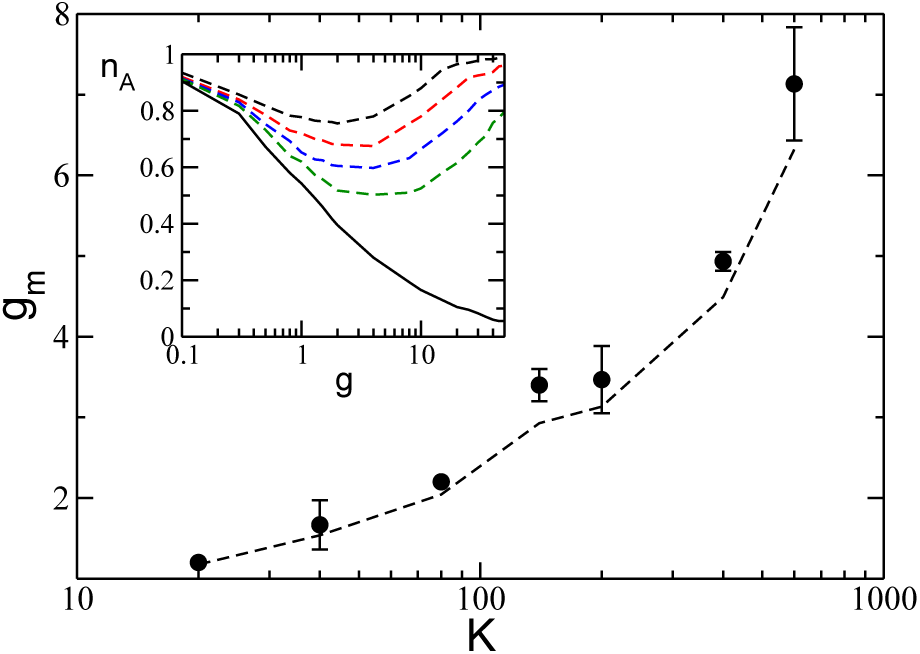
*g*_*m*_ as a function of the in-degree *K.* The symbols refer to numerical data, while the dashed line to the expression (15). Inset: *n*_*A*_ versus *g* for the fully coupled case (solid black line) and for diluted networks (dashed lines), from top to bottom *K* = 20, 40, 80, 140. A network of size *N* = 1400 is evolved during a period *t*_*S*_ = 1 × 10^5^ after discarding a transient of 10^6^ spikes, the data are averaged over 5 different random realizations of the network. Other parameters as in Fig. 1.

The general mechanism behind neurons’ rebirth can be understood by considering the value of the effective neuronal input and of the current fluctuations as a function of *g.* As shown in Fig. 4, the effective input current 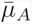, averaged over the active neurons, essentially coincide with the average of the excitability 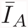 for *g →* 0, where the neurons can be considered as independent one from the others. The inhibition leads to a decrease of 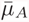, and to a crossing of the threshold *θ* exactly at *g* = *g*_*m*_. This indicates that at *g* < *g*_*m*_ the active neurons, being on average supra-threshold, fire almost tonically inhibiting the losers via a WTA mechanism. In this case the firing neurons are essentially mean-driven and the current fluctuations play a role on the rebirth of silent neurons only on extremely long time scales, this is confirmed by the low values of 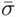 in such a range, as evident from Fig. 4. On the other hand, for *g* > *g*_*m*_, the active neurons are now on average below threshold while the fluctuations dominate the dynamics. In particular, the firing is now extremely irregular being due mainly to reactivation processes. Therefore the origin of the minimum in *n*_*A*_ can be understood as a transition from a mean-driven to a fluctuation-driven regime [67].

A quantitative definition of *g*_*m*_ can be given by requiring that the average input current of the active neurons 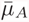 crosses the threshold *θ* at *g* = *g*_*m*_, namely

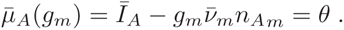

where 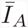 is the average excitability of the firing neurons, while *n*_*Am*_ and 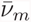 are the number of active neurons and the average frequency at the minimum.

For an uniform distribution *P*(*I*), a simple expression for *g*_*m*_ can be derived, namely

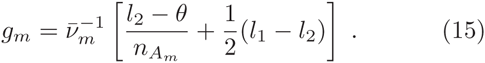

The estimations obtained with this expression are compared with the numerical data reported in Fig. 6 for a network of size *N* = 1,400 and various in-degrees. The overall agreement is more than satisfactory for in-degrees ranging over almost two decades (namely for 20 ≤ *K ≤* 600).

It should be stressed that, as we have verified for various system sizes (namely *N* = 700,1400 and 2800) and for a constant average in-degree *K* = 140, for instantaneous synapses the network is in an heterogeneous asynchronous state for all the considered values of the synaptic coupling. This is demonstrated by the fact that the amplitude of the fluctuations of the average firing activity measured by considering the low-pass filtered linear super-position of all the spikes emitted in the network, vanishes as 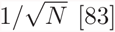 [83]. Therefore, the observed transition at *g* = *g*_*m*_ is not associated to the emergence of irregular collective behaviours as reported for globally coupled heterogeneous inhibitory networks of LIF neurons with delay [42] and of pulse-coupled phase oscillators [80].

## V. EFFECT OF SYNAPTIC FILTERING

In this Section we will investigate how synaptic filtering can influence the previously reported results. In particular, we will consider non instantaneous IPSP with *α*-function profile (3), whose evolution is ruled by a single time scale *τ*_*α*_.

### A. Fully Coupled Networks

Let us first examine the fully coupled topology, in this case we observe analogously to the *δ*-pulse coupling that by increasing the inhibition, the number of active neurons steadily decreases towards a limit where only few neurons (or eventually only one) will survive. At the same time the average frequency also decreases mono tonically as shown in Fig. 7 for two different *τ*_*α*_ differing by almost two orders of magnitude. Furthermore, the mean field estimations (7) and (8) obtained for *n*_*A*_ and 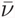 represent a very good approximation also for *α-*pulses (as shown in Fig. 7). In particular, the mean field estimation essentially coincides with the numerical values for slow synapses, as evident from the data reported in Fig. 7 for *τ*_*α*_ = 10 (black filled circles). This can be explained by the fact that for sufficiently slow synapses, with 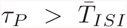, the neurons feel the synaptic input current as continuous, because each input pulse has essentially no time to decay between two firing events. Therefore the mean field approximation for the input current (4) works definitely well in this case. This is particularly true for *τ*_*α*_ = 10, where *τ*_*P*_ = 20 and 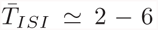 in the range of the considered coupling. While for *τ*_*α*_ = 0.125, we observe some deviation from the mean field results (red squares in Fig. 7) and the reason for these discrepancies reside in the fact that 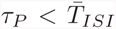 for any coupling strength, therefore the discreteness of the pulses cannot be completely neglected in particular for large amplitudes (large synaptic couplings) analogously to what observed for instantaneous synapses.

**FIG. 7.**
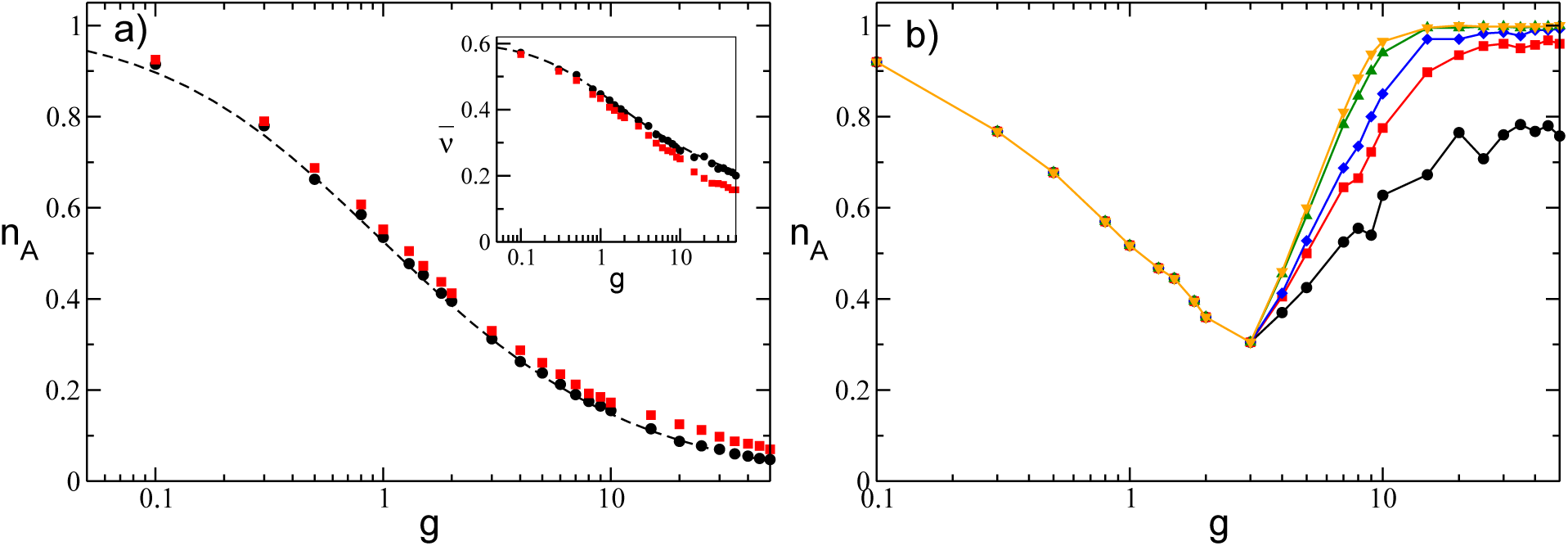
Fraction of active neurons *n*_*A*_ as a function of the inhibition with IPSPs with α-profile for a fully coupled topology (a) and a sparse network (b) with *K* = 20. a) Black (red) symbols correspond to *τ*_*α*_ = 10 (*τ*_*α*_ = 0.125), while the dashed lines are the theoretical predictions (7) and (8) previously reported for instantaneous synapses. The data are averaged over a time window *t*_*s*_ = 1 × 10^5^. Inset: average frequency 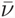 as a function of *g.* b) *n*_*A*_ is measured at successive times : from lower to upper curve the considered times are *t*_*s*_ = {1000, 5000, 10000, 50000, 100000}, while *τ*_*α*_ = 10. The system size is *N* = 400 in both cases, the distribution of excitabilities is uniform with [*l*_1_ : *l*_2_] = [1.0 : 1.5] and *θ* = 1.

### B. Sparse Networks

For the sparse networks *n*_*A*_ has the same qualitative behaviour as a function of the synaptic inhibition observed for instantaneous IPSPs, as shown in Fig. 7 (b) and Fig. 8 (a). The value of *n*_*A*_ decreases with *g* and reaches a minimal value at *g*_*m*_, afterwards it increases towards *n*_*A*_ = 1 at larger coupling. The origin of the minimum of *n*_*A*_ as a function of *g* is the same as for instantaneous synapses, for *g* < *g*_*m*_ the active neurons are subject on average to a supra-threshold effective input 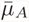, while at larger coupling 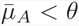, as shown in the inset of Fig. 8 (b). This is true for any value of *τ*_α_, however, this transition from mean- to fluctuation-driven becomes dramatic for slow synapses. As evidenced from the data for the average output firing rate 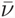 and the average coefficient of variation 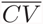, these quantities have almost discontinuous jumps at *g* = *g*_*m*_, as shown in Fig. 9.

**FIG. 8.**
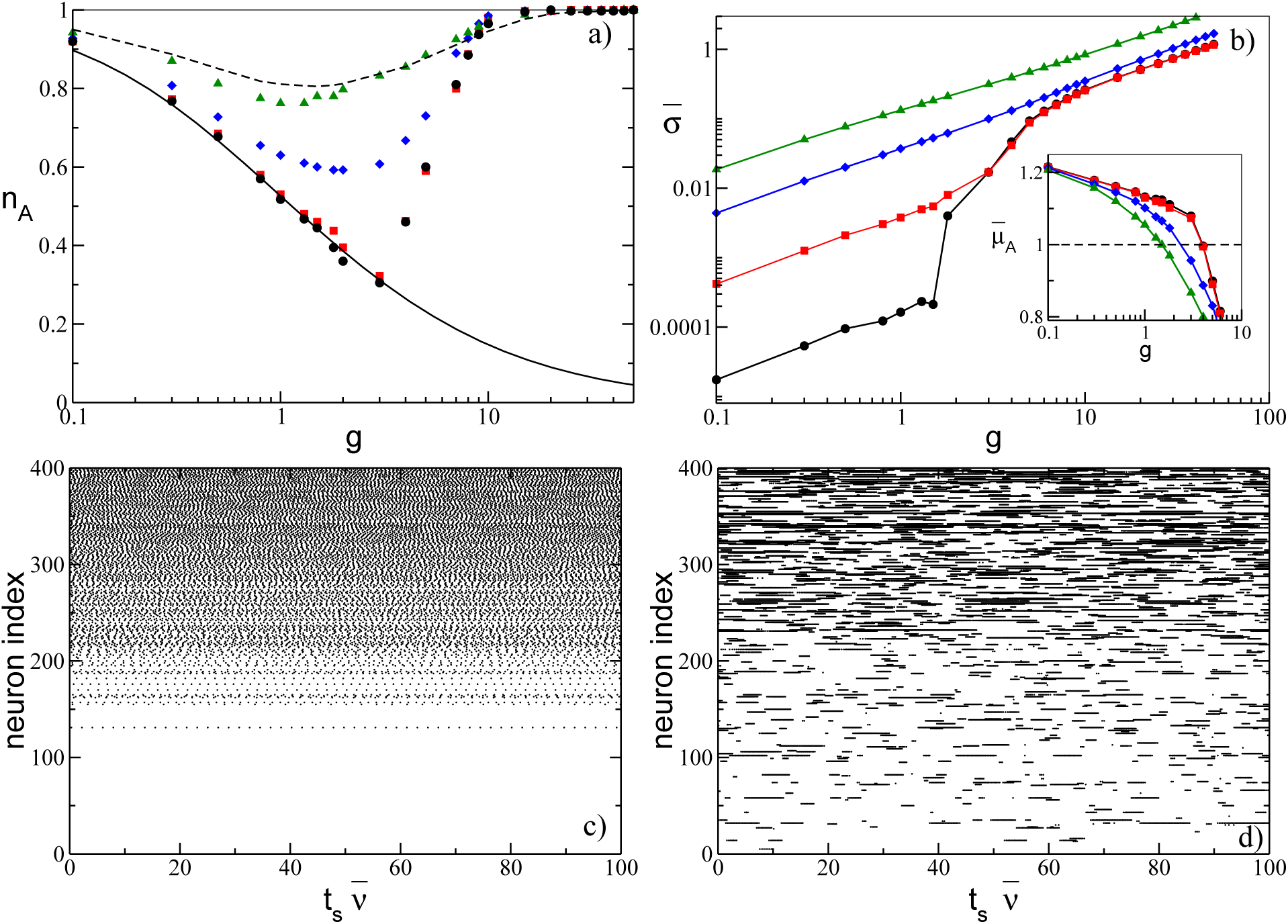
a) Fraction of active neurons for a network of α-pulse coupled neurons as a function of *g* for various *τ*_*α*_: namely, *τ*_*α*_ = 10 (black circles), *τ*_*α*_ = 2 (red squares), *τ*_*α*_ = 0.5 (blue diamonds) and *τ*_*α*_ = 0.125 (green triangles). For instantaneous synapses, the fully coupled analytic solution is reported (solid line), as well as the measured *n*_*A*_ for the sparse network with same level of dilution and estimated over the same time interval (dashed line). b) Average fluctuations of the synaptic current 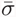 versus *g* for ISPS with *α*-profile the symbols refer to the same *τ*_*α*_ as in panel (a). Inset: Average input current 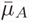 of the active neurons vs *g*, the dashed line is the threshold value *θ* = 1. The simulation time has been fixed to *t*_*s*_ = 1 *×* 10^5^. c-d) Raster plots for two different synaptic strengths for *τ*_*α*_ = 10: namely, c) *g* = 1 corresponds to *n*_*A*_ ≃ 0.52, 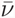 ≃ 0.45 and 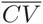 ν 3 × 10^−4^; while i) *g* = 10 to *n*_*A*_ ≃ 0.99, 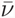 ≃ 0.06 and 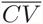 ≃ 4.1. The neurons are ordered according to their intrinsic excitability and the time is rescaled by the average frequency 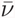. The data have been obtained for a system size *N* = 400 and *K* = 20, other parameters as in Fig. 7.

**FIG. 9.**
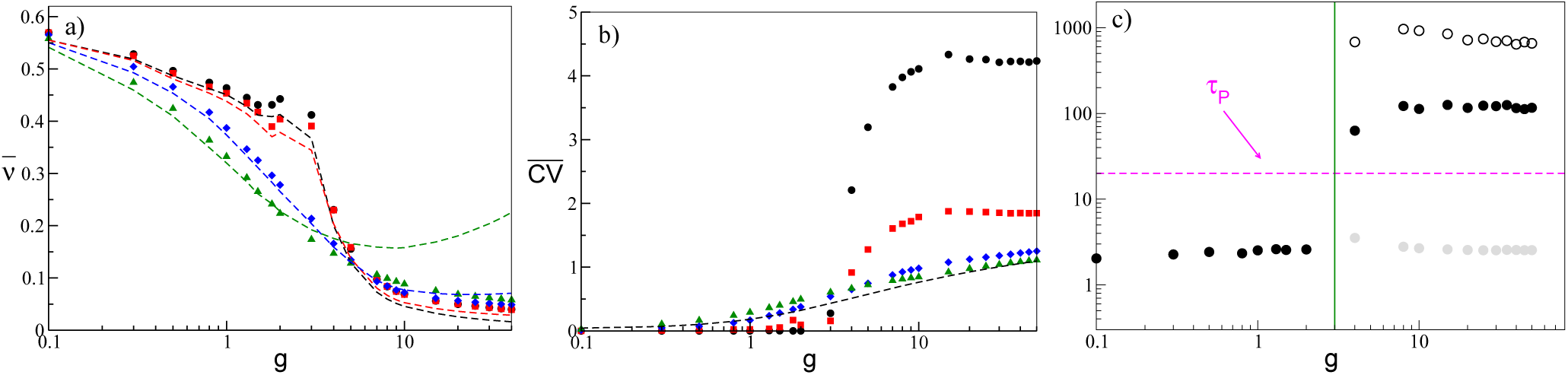
a) Average firing rate 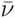 vs *g* for a network of *α*-pulse coupled neurons, for four values of *τ*_*α*_. Theoretical estimations for 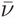 calculated with the adiabatic approach (45) are reported as dashed lines of colors corresponding to the relative symbols. b) Average coefficient of variation 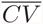 for four values of *τ*_*α*_ as a function of the inhibition. The dashed line refers to the values obtained for instantaneous synapses and a sparse network with the same value of dilution. c): Average inter-spike interval 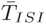 (filled black circles) as a function of *g* for *τ*_*α*_ = 10. For *g* > *g*_*m*_ the average inter-burst interval (empty circles) and the average ISI measured within bursts (gray circles) are also shown, together with the position of *g*_*m*_ (green veritical line). The symbols and colors denote the same *τ*_*α*_ values as in Fig. 8. All the reported data were calculated for a system size *N* = 400 and *K* = 20 and for a fixed simulation time of *t*_*s*_ = 1 × 10^5^.

Therefore, let us first concentrate on slow synapses with *τ*_*α*_ larger than the membrane time constant, which is one for adimensional units. For *g* < *g*_*m*_ the fraction of active neurons is frozen in time, at least on the considered time scales, as revealed by the data in Fig. 7 (b). Furthermore, for *g* < *g*_*m*_, the mean field approximation obtained for the fully coupled case works almost perfectly both for *n*_*A*_ and 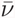, as reported in Fig. 8 (a). The frozen phase is characterized by extremely small values of the current fluctuations 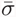 (as shown Fig. 8 (b)) and a quite high firing rate 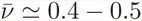 with an associated average coefficient of variation 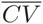 almost zero (see black circles and red squares in Fig. 9). Instead, for *g* > *g*_*m*_ the number of active neurons increases in time similarly to what observed for the instantaneous synapses, while the average frequency becomes extremely small 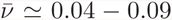 and the value of the coefficient of variation becomes definitely larger than one.

These effects can be explained by the fact that, below *g*_*m*_ the active neurons (the winners) are subject to an effective input 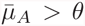 that induces a quite regular firing, as testified by the raster plot displayed in Fig. 8 (c). The supra-threshold activity of the winners joined together with the filtering action of the synapses guarantee that on average each neuron in the network receive an almost continuous current, with small fluctuations in time. These results explain why the mean field approximation still works in the frozen phase, where the fluctuations in the synaptic currents are essentially negligible and unable to induce any neuron’s rebirth, at least on realistic time scales. In this regime the only mechanism in action is the WTA, fluctuations begin to have a role for slow synapses only for *g* > *g*_*m*_. Indeed, as shown in Fig. 8 (b), the synaptic fluctuations 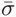 for *τ*_*α*_ = 10 (black circles) are almost negligible for *g* < *g*_*m*_ and show an enormous increase at *g* = *g*_*m*_ of almost two orders of magnitude. Similarly at *τ*_*α*_ = 2 (red square) a noticeable increase of 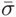 is observable at the transition.

In order to better understand the abrupt changes in 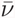 and in 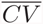 observable for slow synapses at *g* = *g*_*m*_, let us consider the case *τ*_*α*_ = 10. As shown in Fig. 9 (c), 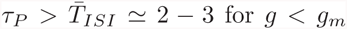, therefore for these couplings the IPSPs have no time to decay between a firing emission and the next one, thus the synaptic fluctuations 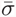 are definitely small in this case, as already shown. At *g*_*m*_ an abrupt jump is observable to large values 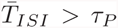, this is due to the fact that now the neurons display bursting activities, as evident from the raster plot shown in Fig. 8 (d). The bursting is due to the fact for *g* > *g*_*m*_ the active neurons are subject to an effective input which is on average sub-threshold, therefore the neurons preferentially tend to be silent. However, due to current fluctuations a neuron can pass the threshold and the silent periods can be interrupted by bursting phases where the neuron fires almost regularly. As a matter of fact, the silent (inter-burst) periods are very long ≃ 700 – 900, if compared to the duration of the bursting periods, namely ≃ 25 – 50, as shown in Fig. 9 (c). This explains the abrupt decrease of the average firing rate reported in Fig. 9 (a). Furthermore, the inter-burst periods are exponentially distributed with a an associated coefficient of variation ≃ 0.8 – 1.0, which clearly indicates the stochastic nature of the switching from the silent to the bursting phase. The firing periods within the bursting phase are instead quite regular, with an associated coefficient of variation ≃ 0.2, and with a duration similar to 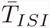 measured in the frozen phase (shaded gray circles in Fig. 9 (c)). Therefore, above *g*_*m*_ the distribution of the ISI exhibits a long exponential tail associated to the bursting activity and this explains the very large values of the measured coefficient of variation. By increasing the coupling the fluctuations in the input current becomes larger thus the fraction of neurons that fires at least once within a certain time interval increases. At the same time, 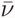, the average inter-burst periods and the firing periods within the bursting phase remain almost constant at *g* > 10, as shown in Fig. 9 (a). This indicates that the decrease of 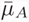 and the increase of 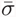 due by the increased inhibitory coupling essentially compensate each other in this range. Indeed, we have verified that for *τ*_*α*_ = 10 and 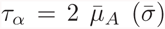 decreases (increases) linearly with *g* with similar slopes, namely 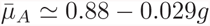 while 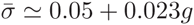.

For faster synapses, the frozen phase is no more present, furthermore due to rebirths induced by current flutuations *n*_*A*_ is always larger than the fully coupled mean field result (8), even at *g* < *g*_*m*_. It is interesting to notice that by decreasing *τ*_*α*_, we are now approaching the instantaneous limit. As indicated by the results reported for *n*_*A*_ in Fig. 8 (a) and 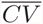 in Fig. 9 (b), in particular for *τ*_*α*_ = 0.125 (green triangles) the data almost collapse on the corresponding values measured for instantaneous synapses in a sparse networks with the same characteristics and over a similar time interval (dashed line). Furthermore, for fast synapses with *τ*_*α*_ < 1 the bursting activity is no more present as it can be appreciated by the fact that at most 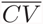 approaches one in the very large coupling limit.

The reported results can be interpreted in the framework of the so-called adiabatic approach developed by Moreno-Bote and Parga in [50, 51], to estimate analytically 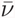. This method applies to LIF neurons with a synaptic time scale definitely longer than the membrane time constant. In these conditions, the output firing rate can be reproduced by assuming that the current fluctuations correspond to colored noise, with a correlation time given by the pulse duration *τ*_*P*_ = 2*τ*_*α*_ (for more details see Appendix D). In this case we are unable to develop an analytic expression for *n*_*A*_, that can be solved self consistently with that for the average frequency. However, once *n*_*A*_ is provided the analytic estimated (45) obtained with the adiabatic approach gives very good agreement with the numerical data for sufficiently slow synapses, namely for *τ*_*P*_ *≥* 1, as shown in Fig. 9(a) for *τ*_*α*_ = 10, 2 and 0.5. The theoretical expression (45) is even able to reproduce the jump in the average frequencies observable at *g*_*m*_ and therefore to capture the bursting phenomenon. By considering *τ*_*P*_ < 1, as expected, the theoretical expression fails to reproduce the numerical data in particular at large coupling (see the dashed green line in Fig. 9(a) corresponding to *τ*_*α*_ = 0.125).

The bursting phenomenon observed for *τ*_*α*_ > 1 and *g* > *g*_*m*_ can be seen at a mean field level as the response of a sub-threshold LIF neuron subject to colored noise with correlation *τ*_*P*_. The neuron is definitely sub-threshold, but in presence of a large fluctuation it can be lead to fire and due to the finite correlation time it can stays suprathreshold regularly firing for a period ≃ *τ*_*P*_. Indeed, the average bursting periods we measured are of the order of *τ*_*P*_= 2*τ*_*α*_, namely ≃ 27 − 50 (≃ 7 − 14) for *τ*_*α*_ = 10 (*τ*_*α*_ = 2)

As a final point, to better understand the dynamical origin of the measured fluctuations in this deterministic model, we have estimated the maximal Lyapunov exponent **λ**. As expected from previous analysis, for non-instantaneous synapses we can observe the emergence of regular chaos in purely inhibitory networks [1, 31, 87]. In particular, for sufficiently fast synapses, we typically note a transition from a chaotic state at low coupling to a linearly stable regime (with **λ** < 0) at large synaptic strengths, as shown in Fig. 10 (a) for *τ*_*α*_ = 0.125. Despite the fact that current fluctuations are monotonically increasing with the synaptic strength. Therefore, fluctuations are due to chaos, at small coupling, while at larger *g* they are due to finite amplitude instabilities, as expected for stable chaotic systems [3]. However, the passage from positive to negative values of the maximal Lyapunov exponent is not related to the transition occurring at *g*_*m*_ from a mean-driven to a fluctuation-driven dynamics in the network.

**FIG. 10.**
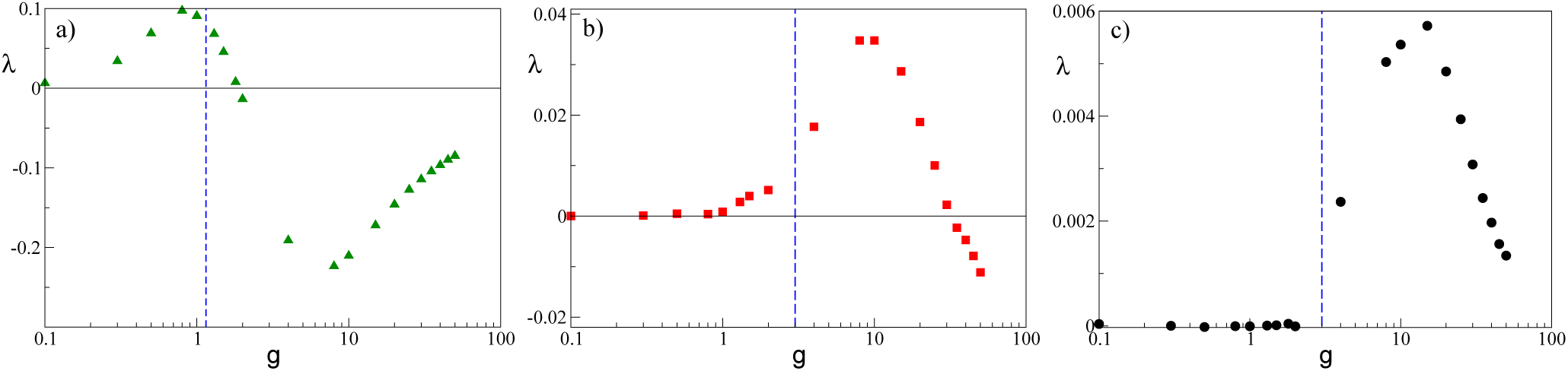
Maximal Lyapunov exponent *λ* versus *g* for a network of *α*-pulse coupled neurons, for *τ*_*α*_ = 0.125 (a), *τ*_*α*_ = 2 (b) and *τ*_*α*_ = 10 (c). The blue dashed vertical line denote the *g*_*m*_ value. All the reported data were calculated for a system size *N* = 400 and *K* = 20 and for simulation times 5 × 10^4^ ≤ *t*_*s*_ ≤ 7 × 10^5^ ensuring a good convergence of *λ* to its asymptotic value. The other parameters are as in in Fig. 7.

For slow synapses, **λ** is essentially zero at small coupling in the frozen phase characterized by tonic spiking of the neurons, while it becomes positive by approaching *g*_*m*_. For larger synaptic strengths **λ**, after reaching a maximal value, decreases and it can become eventually negative at *g* >> *g*_*m*_, as reported in Fig. 10 (b-c). Only for extremely slow synapses, as shown in Fig. 10 (c) for *τ*_*α*_ = 10, the chaos onset seems to coincide with the transition occurring at *g*_*m*_. These findings are consistent with recent results concerning the emergence of asynchronous rate chaos in homogeneous inhibitory LIF networks with deterministic [27] and stochastic [32] evolution. However, a detailed analysis of this aspect goes beyond the scope of the present paper.

## VI. DISCUSSION

In this paper we have shown that the effect reported in [1, 65] is observable whenever two sources of quenched disorder are present in the network: namely, a random distribution of the neural properties and a random topology. In particular, we have shown that neuron’s death due to synaptic inhibition is observable only for heterogeneous distributions of the neural excitabilities. Furthermore, in a globally coupled network the less excitable neurons are silenced for increasing synaptic strength until only one or few neurons remain active. This scenario corresponds to the winner-takes-all mechanism via lateral inhibition, which has been often invoked in neuroscience to explain several brain functions [86]. WTA mechanisms have been proposed to model hippocampal CA1 activity [16], as well as to be at the basis of visual velocity estimate [25], and to be essential for controlling visual attention [29].

However, most brain circuits are characterized by sparse connectivity [10, 39, 59], in these networks we have shown that an increase in inhibition can lead from a phase dominated by neuronal death to a regime where neuronal rebirths take place. Therefore the growth of inhibition can have the counter-intuitive effect to activate silent neurons due to the enhancement of current fluctuations. The reported transition is characterized by a passage from a regime dominated by the almost tonic activity of a group of neurons, to a phase where sub-threshold fluctuations are at the origin of the irregular firing of large part of the neurons in the network. For instantaneous synapses, the average first and second moment of the firing distributions have been obtained together with the fraction of active neurons within a mean-field approach, where the neuronal firing is interpreted as an activation process driven by synaptic shot noise [69].

For finite synaptic time smaller than the characteristic membrane time constant the scenario is similar to the one observed for instantaneous synapses. However, the transition from mean-driven to fluctuation-driven dynamics becomes dramatic for sufficiently slow synapses. In this situation one observes for low synaptic strength a frozen phase, where the synaptic filtering washes out the current fluctuations leading to an extremely regular dynamics controlled only by a WTA mechanism. As soon as the inhibition is sufficiently strong to lead the active neurons below threshold the neuronal activity becomes extremely irregular exhibiting long silent phases interrupted by bursting events. The origin of these bursting periods can be understood in terms of the emergence of correlations in the current fluctuations [50].

In our model, the random dilution of the network connectivity is a fundamental ingredient to generate current fluctuations, whose amplitude is controlled by the average network in-degree *K*. A natural question is if the reported scenario will be still observable in the thermodynamic limit. On the basis of previous studies we can affirm that this depends on how *K* scales with the system size [23, 41, 74]. In particular, if *K* stays finite for *N →* ∞ the transition will still be observable, while for *K* diverging with *N* the fluctuations will become negligible for sufficiently large system sizes not allowing for neuronal rebirths and the dynamics will be controlled only by the WTA mechanism.

As a further aspect, we show that the considered model is not chaotic for instantaneous synapses, in such a case we observe irregular asynchronous states due to stable chaos [62]. The system can become truly chaotic only for finite synaptic times [3, 31], however we clearly report clear indications that for synapses faster than the membrane time constant *τ*_*m*_ the passage from mean-driven to fluctuation-driven dynamics is not related to the onset of chaos. Only for extremely slow synapses we have numerical evidences that the appearance of the bursting regime could be related to a passage from a zero Lyapunov exponent to a positive one, in agreement with the results reported in [27, 32] for homogeneous inhibitory networks. These preliminary indications demand for future more detailed investigations of deterministic spiking networks in order to relate fluctuation-driven regime and chaos onsets. Furthermore, we expect that it will be hard to distinguish whether the erratic current fluctuations are due to regular chaos or stable chaos on the basis of the analysis of the network activity, as also pointed out in [31].

For what concerns the biological relevance of the presented model, we can attempt a comparison with experimental data obtained for MSNs in the striatum. This population of neurons is fully inhibitory with lateral connections, which are sparse (connection probability ≃ 10 – 20% [75, 79]), unidirectional and relatively weak [76]. The dynamics of these neurons in behaving mice reveal a low average firing rate with irregular firing activity (bursting) with associated large coefficient of variation [47]. As we have shown, these features can be reproduced by a sparse networks of LIF neurons with sufficiently slow synapses at *g* > *g*_*m*_ and *τ*_*α*_ > *τ*_*m*_. For values of the membrane time constant which are comparable to the ones measured for MSNs [58, 60] (namely *τ*_*m*_ ≃ 10 – 20 msec), the model is able to capture even quantitatively some of the main aspects of the MSNs dynamics, as shown in Table I. We obtain a reasonable agreement with the experiments for sufficiently slow synapses, while the interaction among MSNs is mainly mediated by GABA_*A*_ receptors, which are characterized by IPSP durations of the order of ≃ 5 – 20 ms [35, 79]. However, apart the burst duration, which is definitely shorter, all the other aspects of the MSN dynamics can be already captured for *τ*_*α*_ = 2*τ*_*m*_ (with *τ*_*m*_ = 10 ms) as shown in Table I. Therefore, we can safely affirm, as also suggested in [65], that the fluctuation driven regime emerging at *g* > *g*_*m*_ is the most appropriate in order to reproduce the dynamical evolution of this population of neurons.

Other inhibitory populations are present in the basal ganglia, in particular two coexisting inhibitory populations, *arkypallidal* (Arkys) and *prototypical* (Protos) neurons, have been recently discovered in the external globus pallidus [43]. These populations have distinct physiological and dynamical characteristics and have been shown to be fundamental for action suppression during the performance of behavioural tasks in rodents [44]. Protos are characterized by a high firing rate ≃ 47 Hz and a not too large coefficient of variation (namely, *CV* ≃ 0.58) both in awake and slow wave sleep (SWS) states, while Arkys have a clear bursting dynamics with *CV* ≃ 1.9 [18, 44]. Furthermore, the firing rate of Arkys is definitely larger in the awake state (namely, ≃ 9 Hz) with respect to the SWS state, where the firing rates are ≃ 3 − 5 Hz [44].

On the basis of our results, Protos can be modeled as LIF neurons with reasonable fast synapses in a mean driven regime, namely with a synaptic coupling *g* < *g*_*m*_, while Arkys should be characterized by IPSP with definitely longer duration and they should be in a fluctuation driven phase as suggested from the results reported in Fig. 9. Since, as shown in Fig. 9 (a), the firing rate of inhibitory neurons decrease by increasing the synaptic strenght *g* we expect that the passage from awake to slow wave sleep should be characterized by a reinforcement of Arkys synapses. Our conjectures about Arkys and Protos synaptic properties based on their dynamical behaviours ask for for experimental verification, which we hope will happen shortly.

**TABLE I.** Comparison between the results obtained for slow *α-* synapses and experimental data for MSNs.The numerical data refer to results obtained in the bursting phase, namely for synaptic strenght *g* in the range [10 : 50], for simulation times *t*_*s*_ = 1 × 10^5^, *N* = 400 and *K* = 20. The experimental data refer to MSNs population in striatum for free behaving wild type mice, data taken from [47].

| $\tau_\alpha/\tau_m$ | $\tau_m$ (msec) | Spike Rate (Hz) | $\overline{CV}$ | Burst Duration (msec) | Spike Rate within Bursts (Hz) |
| --- | --- | --- | --- | --- | --- |
| 2 | 10 | 4-6 | $\simeq 1.8$ | $100 \pm 40$ | $42 \pm 2$ |
| | 20 | 2-3 | $\simeq 1.8$ | $200 \pm 80$ | $21 \pm 1$ |
| 10 | 10 | 4-6 | $\simeq 4.2$ | $400 \pm 150$ | $41 \pm 2$ |
| | 20 | 2-3 | $\simeq 4.2$ | $800 \pm 300$ | $20 \pm 1$ |
| | Experimental data | 2-3 | $\simeq 1.5 - 3$ | $500 - 1100$ | $31 \pm 15$ |

Besides the straightforward applicability of our findings to networks of pulse-coupled oscillators [48], it has been recently shown that LIF networks with instantaneous and non-instantaneous synapses can be transformed into the Kuramoto-Daido model [17, 36, 61]. Therefore, we expect that our findings should extend to phase oscillator arrays with repulsive coupling [77]. This will allow for a wider applicability of our results, due to the relevance of limit-cycle oscillators not only for modeling biological systems [84], but also for the many scientific and technological applications [19, 57, 70, 73].

## ACKNOWLEDGMENTS

Some preliminary analysis on the shot noise has been performed in collaboration with A. Imparato, the complete results will be reported elsewhere [54]. We thank for useful discussions B. Lindner, G. Mato, G. Giacomelli, S. Gupta, A. Politi, MJE Richardson, R. Schmidt, M.Timme. This work has been partially supported by the European Commission under the program “Marie Curie Network for Initial Training”, through the project N. 289146, “Neural Engineering Transformative Technologies (NETT)” (D.A.-G., S.O., and A.T), by the A*MIDEX grant (No. ANR-11-IDEX-0001-02) funded by the French Government “program Investissements d’Avenir” (D.A.-G. and A.T.), and by “Departamento Adminsitrativo de Ciencia Tecnologia e Innovacion - Colciencias” through the program “Doctorados en el exterior - 2013” (D.A.-G.). This work has been completed at Max Planck Institute for the Physics of Complex Systems in Dresden (Germany) as part of the activity of the Advanced Study Group 2016 entitled “From Microscopic to Collective Dynamics in Neural Circuits”.

## APPENDIX A EVENT DRIVEN MAPS

By following [53, 89] the ordinary differential equations (1) and (2) describing the evolution of the membrane potential of the neurons can be rewritten exactly as discrete time maps connecting successive firing events occurring in the network. In the following we will report explicitly such *event driven maps* for the case of instantaneous and *α* synapses.

For instantaneous PSPs, the event-driven map for neuron *i* takes the following expression:

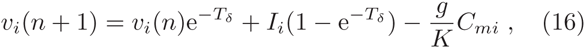

where the sequence of firing times {*t*_*n*_} in the network is denoted by the integer indices {*n*}, m is the index of the neuron firing at time *t*_*n*+1_ and *T*_*δ*_ ≡ *t*_*n*+1_ – *t*_*n*_ is the inter-spike interval associated with two successive neuronal firing. This latter quantity is calculated from the following expression:

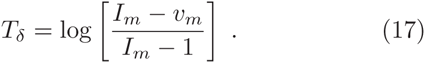

For *α*-pulses, the evolution of the synaptic current *E*_*i*_(*t*), stimulating the i-th neuron can be expressed in terms of a second order differential equation, namely

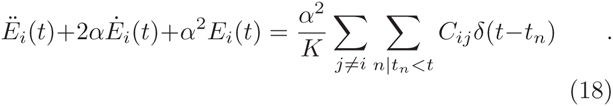

Eq. (18) can be rewritten as two first order differential equations by introducing the auxiliary variable *Q* ≡ *Ė*_*i*_ – α*E*_*i*_, namely

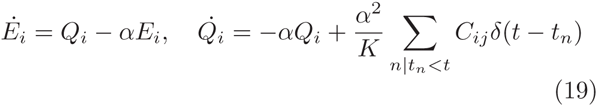

Finally, the equations (1) and (19) can be exactly integrated from the time *t*_*n*_, just after the deliver of the *n*-th pulse, to time *t*_*n*+1_ corresponding to the emission of the (*n* + 1)-th spike, to obtain the following event driven map:

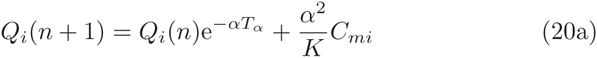

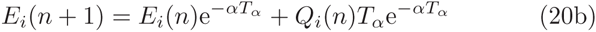

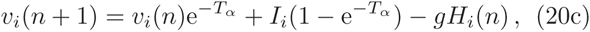

In this case, the inter-spike interval *T*_*α*_ ≡ *t*_*n*+1_ – *t*_*n*_ should be estimated by solving self-consistently the following expression

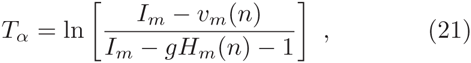

where the explicit expression for *H*_*i*_(*n*) appearing in equations (20c) and (21) is

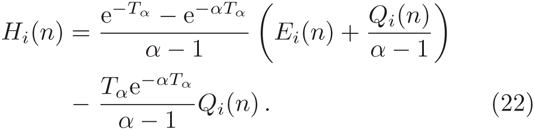

The model so far introduced contains only adimensional units, however, the evolution equation for the membrane potential (1) can be easily re-expressed in terms of dimensional variables as follows

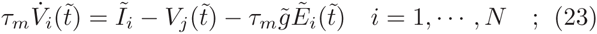

where we have chosen *τ*_*m*_ = 10 ms as the membrane time constant, 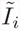 represents the neural excitability and the external stimulations. Furthermore, 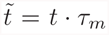, the field 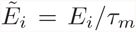 has the dimensionality of a frequency and 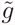 of a voltage. The currents 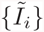 have also the dimensionality of a voltage, since they include the membrane resistance.

For the other parameters/variables the transformation to physical units is simply given by

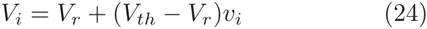

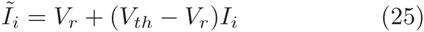

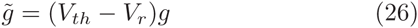

where *V*_*r*_ = −60 mV and *V*_*th*_ = −50 mV are realistic values of the membrane reset and threshold potential. The isolated *i*-th LIF neuron is supra-threshold whenever 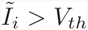.

## APPENDIX B AVERAGE FIRING RATE FOR INHIBITORY SHOT NOISE

In this Appendix, by following the approach in [69] we derive the average firing rate of a supra-threshold LIF neuron subject to inhibitory synaptic shot noise of constant amplitude *G*, namely

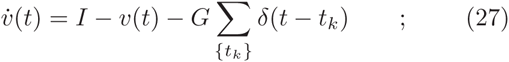

where *I* > 1. The synaptic pulses reach the neuron at Poisson-distributed arrival times with a rate *R.* In order to find the firing rate response of the LIF neuron we introduce the probability density *P*(*v*) and the flux *J*(*v*) associated to the membrane potentials, these satisfy the continuity equation:

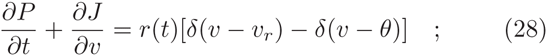

where *r*(*t*) is the instantaneous firing rate. The flux can be decomposed in an average drift term plus the inhibitory part, namely

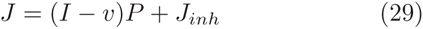

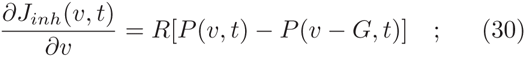

The set of equations (28) to (30) is complemented by the boundary conditions:

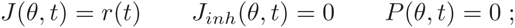

and by the requirement that membrane potential distribution should be normalized, i.e

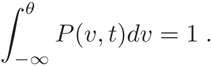

By introducing bilateral Laplace transforms 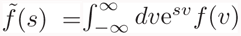 and by performing some algebra along the lines described in [69] it is possible to derive the analytic expression for the average firing rate

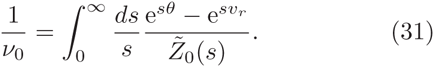

where 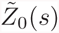 is the Laplace transform of the sub-threshold probability density. Namely, it reads as

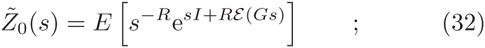

where 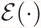 is the exponential integral, *E* = e^−*R*(Γ+lnG)^ is the normalization constant ensuring that the distribution *Z*_0_(*v*) is properly normalized, and Γ is the Euler-Mascheroni constant.

## APPENDIX C COEFFICIENT OF VARIATION FOR INHIBITORY SHOT NOISE

In order to derive the coefficient of variation for the shot noise case it is necessary to obtain the first two moments of the first-passage-time density *q*(*t*). By following the same approach as in Appendix B, the continuity equation associated to the response to a spike given at time *t* = 0 is the following

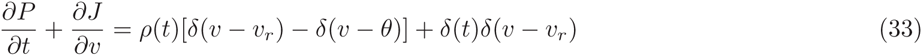

where *ρ*(*t*) is the spike-triggered rate. As suggested in [69], Eq. (33) can be solved by performing a Fourier transform in time and a bilateral Laplace transform in membrane potential. This allows to obtain the Fourier transform of the spike-triggered rate, namely

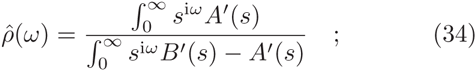

where 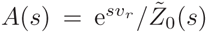 and 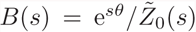. The Fourier transform of the first-passage-time density is 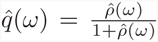 and the first and second moment of the distribution are given by

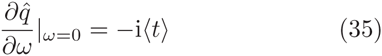

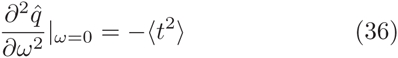

The integrals appearing in Eq. (34) cannot be exactly solved, therefore we have expanded it to the second order obtaining

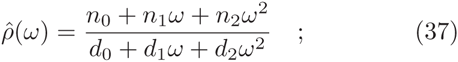

where *n*_0_ = −1 and *d*_1_ = −i*n*_0_/*ν*_0_ and

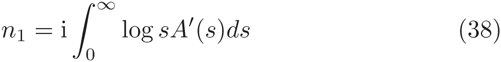

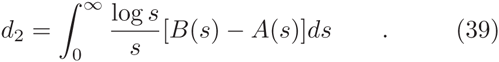

From the expression (34) we can finally obtain the first and second moment of *q*(*t*), namely

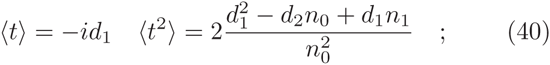

Once these quantities are known the coefficient of variation can be easily estimated for each neuron with excitability *I.*

## APPENDIX D AVERAGE FIRING RATE FOR SLOW SYNAPSES

In Section V we have examined the average activity of the network with synaptic transmission described by *α-*functions. In presence of synaptic filtering and when the synaptic time constant is larger than the membrane time constant one can apply the so-called adiabatic approach to derive the firing rate of a single neuron, as reported in [50, 51].

In this approximation, the output firing rate *ν*_0_ of the single neuron driven by a slowly varying stochastic input current *z* with an arbitrary distribution *P*(*z*) is given by

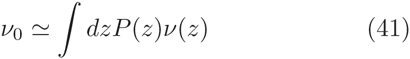

where *ν*(*z*) is the input to rate transfer function of the neuron under a stationary input which for the LIF neuron is simply:

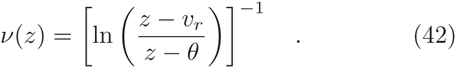

In our network model, a single neuron receives an average current *µ* given by Eq. (4) with a standard deviation *σ* given by Eq. (10).

The synaptic filtering induces temporal correlations in the input current *z*, which can be written as:

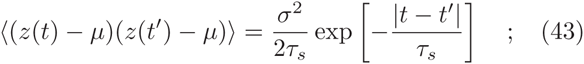

here *τ*_*s*_ is the synaptic correlation time. In the case of *α-*pulses, where the rise and decay synaptic times coincide, we can approximate the correlation time as *τ*_*s*_ = *τ*_*P*_ = 2*τ*_*α*_.

Analogously to the diffusion approximation [8, 9, 14], the input currents are approximated as a Gaussian noise with mean *µ* and variance 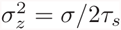.

Therefore, the single neuron output firing rate reads as

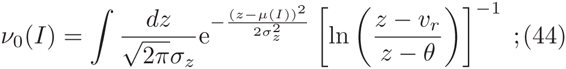

where *I* is the neuronal excitability.

The average firing rate of the LIF neurons in the network, characterized by an excitability distribution *P*(*I*), can be estimated as

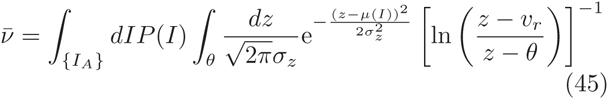

where we impose the self-consistent condition that the average output frequency is equal to the average input one.

